# Lipid Composition Controls the Huntingtin Exon1 Membrane-Association and Differentially Modulates its Flanking Regions Dynamics

**DOI:** 10.1101/2025.09.30.679536

**Authors:** Tânia Sousa, Gonçalo Damas, Ana Coutinho, Nuno Bernardes, Ana Azevedo, Manuel Prieto, Ana M. Melo

## Abstract

The pathological expansion of the polyglutamine (polyQ) repeat within the first exon of huntingtin (Httex1) protein is a defining hallmark of Huntington’s disease (HD). Multiple evidence supports that the membrane recruitment of Httex1 is critical for its self-assembly and related toxicity in HD. In this work, we quantitatively examined the early steps of monomeric Httex1(23Q) association with lipid membranes and its impact on the conformational dynamics of the adjacent polyQ regions - the N-terminal N17 segment and C-terminal proline-rich region (PRR). A broad range of membrane physical properties was explored, including zwitterionic and anionic lipids, and also co-existing liquid-ordered and liquid-disordered phases. Two single cysteine mutants were engineered at the N- and C-termini of Httex1(23Q) and fluorescently-labeled with acrylodan or Atto 488 to probe their local polarity and flexibility, respectively. Our results indicate that Httex1- 23Q preferentially binds to negatively-charged lipid vesicles, and to a lower extent to liquid ordered/disordered phases. The N-terminal N17 segment inserts deeply into anionic membranes, adopting a less flexible state than in aqueous solution. At variance, the C-terminal PRR remains highly dynamic and solvent exposed in the Httex1-23Q membrane-bound state, preserving its intrinsic disordered features across all lipid compositions used. Altogether, our work provides unique insight into the distinct roles of each flanking polyQ region in mediating httex1-lipid binding, and how the lipid composition further modulates these early interaction steps.

## INTRODUCTION

Huntington’s disease (HD) is a progressive inherited neurodegenerative disorder caused by a CAG trinucleotide repeat expansion (> 36 CAG), encoding for an extended polyglutamine (polyQ) stretch near the N-terminus of huntingtin (Htt) protein (MacDonald et al. 1993). Mutant Htt exhibits a high propensity to self-assemble into oligomers /amyloid-like aggregates and forms intracellular inclusion bodies (DiFiglia et al. 1997; Iuchi et al. 2003), which are ultimately implicated in neuronal dysfunction and toxicity. Most biophysical and biochemistry studies have been mainly focused on the highly toxic Htt fragment spanning exon 1 (Httex1), that results from aberrant splicing and proteolysis of the mutant Htt protein (Wellington et al. 2002; Sathasivam et al. 2013). This Httex1 fragment is sufficient to replicate much of HD’s pathology and progression (Mangiarini et al. 1996), serving as both biomarker and promising therapeutic target in HD.

Httex1 comprises a central polyQ domain flanked by a 17 amino acid N-terminal sequence (N17) and a C-terminal proline-rich region (PRR) (Figure 1a) (Wetzel 2012; Adegbuyiro et al. 2017). In the monomeric solution state, Httex1 exhibits disordered features (as an intrinsically disordered protein (IDP)). Recent experimental evidence obtained from nuclear magnetic resonance (NMR) (Newcombe et al. 2018) and single-molecule Förster resonance energy transfer (smFRET) measurements (Warner et al. 2017) suggests it adopts a “tadpole-like” conformation. In this ensemble, N17/polyQ forms a collapsed head and the PRR extends as a semi-flexible tail (Warner et al. 2017; Newcombe et al. 2018). While the Httex1 aggregation is strongly correlated with the polyQ-length, both flanking regions of the polyQ tract - N17 and PRR - can also control its aggregation behavior in solution (Thakur et al. 2009; Shen et al. 2016) and in the presence of biological membranes (Burke et al. 2013b).

**Figure 1:**
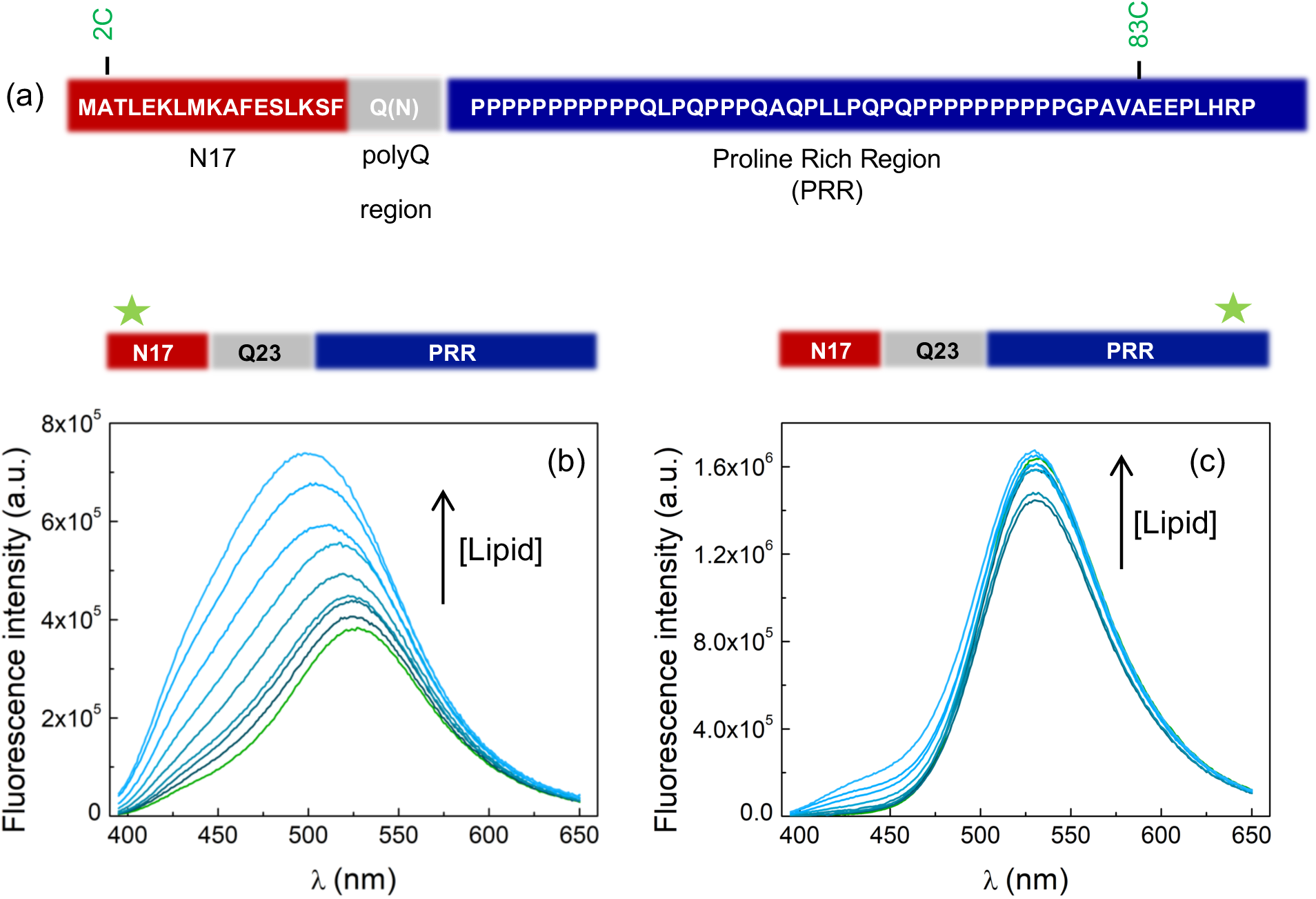
Designed acrylodan-labeled Httex1(23Q) variants for probing membrane binding. (a) Primary sequence of Httex1 and schematic representation of its domains with the N-terminal N17 segment in red, polyQ in grey and C-terminal PRR in blue. The engineered single-cysteine mutants (2C and 83C) for fluorophore attachment are indicated in green. (b-c) Fluorescence emission spectra of 0.6 µM (b) HttAcrylodan-2C and (c) HttAcrylodan--83C in buffer (50 mM HEPES, 50mM NaCl, pH 7.4) and with increasing concentrations of pure POPC LUVs (ranging from 2.5µM to 1mM).

Accumulated evidence supports that biological membranes are key players for both Htt function and in disease context (DiFiglia et al. 1995; Velier et al. 1998; Rockabrand et al. 2007). In its functional state, Htt is widely localized in the cytoplasm (DiFiglia et al. 1995) and is actively involved in a range of membrane-associated processes, including vesicular trafficking and synaptic transmission (DiFiglia et al. 1995; Velier et al. 1998; Caviston and Holzbaur 2009; Vitet et al. 2023). Both Htt and N-terminal fragments closely associate with multiple membranous organelles of diverse lipid compositions (such as endoplasmic reticulum (ER) (Atwal et al. 2007; Rockabrand et al. 2007; Ueda et al. 2016), mitochondria (Panov et al. 2002; Choo et al. 2004) and Golgi apparatus (DiFiglia et al. 1995; Velier et al. 1998; Rockabrand et al. 2007)), and synaptic vesicles (Li et al. 2000). Moreover, pathogenic Httex1 inclusions can sequester lipids that ultimately compromise organelle integrity (Bäuerlein et al. 2017; Riguet et al. 2021). Multiple *in vitro* biophysical studies have shown that lipid bilayers strongly trigger Httex1 aggregation and fibrillation (Burke et al. 2013b; Pandey et al. 2018; Beasley et al. 2021). Likely, the increase of Httex1 local concentration at membrane surface may create nucleation sites or stabilize toxic conformations (as proposed for other aggregation-prone proteins (Butterfield and Lashuel 2010)). Such aberrant Httex1 aggregation can also lead to membrane disruption and permeabilization (Burke et al. 2013a; Bäuerlein et al. 2017). Furthermore, changes in cholesterol and lipid metabolism have been consistently reported in HD (Valenza et al. 2005; Leoni and Caccia 2014; Aditi et al. 2016; Di Pardo et al. 2017), suggesting that lipid membranes are integral components of HD pathology. The membrane recruitment of Httex1 is directly mediated by the N17 segment, which undergoes a transition from a disordered state in solution to an amphipathic α-helix upon membrane anchoring (Atwal et al. 2007; Michalek et al. 2013; Adegbuyiro et al. 2017; Tao et al. 2019). This disorder-to-order conformational transition is a shared feature with other aggregation-prone proteins/peptides (such as a-synuclein, Ab-peptide and islet amyloid peptide (Butterfield and Lashuel 2010)). The N17 lipid-binding motif is also a hotspot for post-translational modifications (PTMs) that critically influence Httex1 aggregation propensity (DeGuire et al. 2018; Sedighi et al. 2020; Chiki et al. 2021; Gottlieb et al. 2021) and also control membrane interaction (Chaibva et al. 2016; DeGuire et al. 2018; Tao et al. 2019; Groover et al. 2020; Sedighi et al. 2020; Adegbuyiro et al. 2022). Specifically, both acetylation at lysines (Lys) 6, 9 and 15 (Chaibva et al. 2016) and phosphomimetic mutations at serines (Ser) 13 and 16 (Tao et al. 2019) were reported to reduce membrane binding.

Considerable effort has been devoted for characterizing the complex aggregation mechanism of Httex1 at lipid membranes. However, the elucidation of the cumulative effects of distinct membrane physical properties on Httex1 membrane partitioning remains incomplete. Multiple Httex1 aggregation and membrane-permeabilization studies have used so far total brain lipid extracts (TBLE from Avanti), where ∼ 58% are of unknow composition, challenging the evaluation of specific lipid effects. For this complex lipid mixture, Httex1 fibrillation was reduced compared with solution (Beasley et al. 2019). In contrast, anionic vesicles (containing 1-palmitoyl-2-oleoyl- *sn*-glycero-3-phosphoserine (POPS), or 1-palmitoyl-2-oleoyl-*sn*-glycero-3-phosphoglycerol (POPG)) significantly accelerated Httex1 aggregation at low lipid:protein ratios (Beasley et al. 2021). Moreover, the impact of membrane phase separation remains unknown as the effects of cholesterol (Chol) and sphingomyelin (SM) have been mostly evaluated in the context of TLBE mixture. Here, cholesterol or sphingomyelin were further shown to modify Httex1 aggregation and membrane insertion (Gao et al. 2016; Chaibva et al. 2018). These previous biophysical studies were mainly performed under high-prone aggregation conditions (typically with Httex1 concentrations at 𝜇M range) and investigated limited lipid compositions.

Here, we employed a systematic approach to quantify the initial interaction steps of monomeric Httex1 with lipid membranes and its effects on N17 and PRR conformational dynamics. Two single-cysteine (Cys) mutants were engineered at N- and C-termini of Httex1(23Q) and further labeled with acrylodan to evaluate their solvent exposure (membrane interfacing), or with Atto- 488 maleimide to analyze their rotational motions. We explored a wide range of membrane physical properties, including pure fluid, negatively charged surfaces, and with co-existing liquid ordered-disordered phases. Our work provides new insights into the complex interplay of surface charge and phase separation on controlling the initial membrane association.

## RESULTS

### The N- and C-termini of Httex1(23Q) experience different local polarities in solution

All experiments were performed with a tag-free Httex1(23Q) produced under native conditions (without organic solvents) using a SUMO Fusion Strategy (based on the Champion™ pET SUMO Expression System, Invitrogen) as described in Material and Methods. This Httex1(23Q) variant enables to probe the monomeric Httex1 state and avoids its extensive oligomerization / aggregation. We took advantage that Httex1 does not contain any natural Cys and introduced a single-Cys mutation at position A2C (N-terminal N17 segment) or A83C (C-terminal PRR) (Figure 1a) in Httex1(23Q). These were first site-specifically labeled with the fluorophore acrylodan to directly map the local polarity of each flanking polyQ region in solution and in its membrane-bound state. Specifically, the fluorescence emission properties of acrylodan are highly sensitive to changes in the polarity of its local environment; with a red-shifted emission indicating a more polar solvent and a blue-shifted emission revealing exposure to a hydrophobic medium (Prendergast et al. 1983). These acrylodan-labeled Httex1(23Q) -2C and -83C constructs (hereafter referred to as Httex1Acrylodan-2C and Httex1Acrylodan-83C, respectively) were initially characterized in aqueous solution through steady-state and picosecond time-resolved fluorescence experiments.

Both acrylodan-labeled Cys variants exhibited a red-shifted fluorescence emission in solution (in the range of the fluorescence emission maximum of acrylodan in water of 540 nm) (Prendergast et al. 1983), reflecting that both N- and C- termini of Httex1(23Q) are largely solvent-exposed as typically expected for disordered proteins (Arya et al. 2018; Chowdhury et al. 2019). However, the fluorescence emission spectrum of acrylodan coupled to residue 2C was slightly broader compared to Httex1Acrylodan-83C and accordingly displayed a shorter fluorescence spectral center-of-mass, 〈𝜆〉 (〈𝜆〉*_!"#$%&&’(_* = 526 nm and 〈𝜆〉*_)*"#$%&&’(_*= 543 nm in aqueous solution) (Figure 1b,c and Table 1). These data are consistent with Httex1Acrylodan-2C experiencing a slightly less polar and more heterogenous microenvironment, which may indicate that the N17 segment adopts transient secondary structures or a subtle more collapsed ensemble in solution. Moreover, the amplitude-weighted mean fluorescence lifetime, 〈𝜏〉, for acrylodan covalently-linked at 2C was shorter than for 83C in buffer (〈𝜏〉*_!"#$%&&’(_*= 0.97 ns and 〈𝜏〉*_)*"# $%&&’(_*= 1.27 ns) (Table 1). Only two lifetime components were required to describe the fluorescence intensity decays of Httex1Acrylodan-83C (typical components: 𝜏,=0.72 ± 0.01 ns and 𝜏*_!_*=2.07 ± 0.02 ns). Instead, the fluorescence intensity decays for Httex1Acrylodan-2C were more complex and their analysis required fitting four lifetime components (typical components: 𝜏,= 0.11 ± 0.01 ns, 𝜏*_!_*= 0.51 ± 0.05 ns, 𝜏*_*_*= 1.61 ± 0.24 ns and 𝜏*_-_*=3.85 ± 0.44 ns), which indicates an overall more heterogeneous microenvironment within the N17 domain (probably with intramolecular quenchers within a more collapsed ensemble).

**Table 1.**
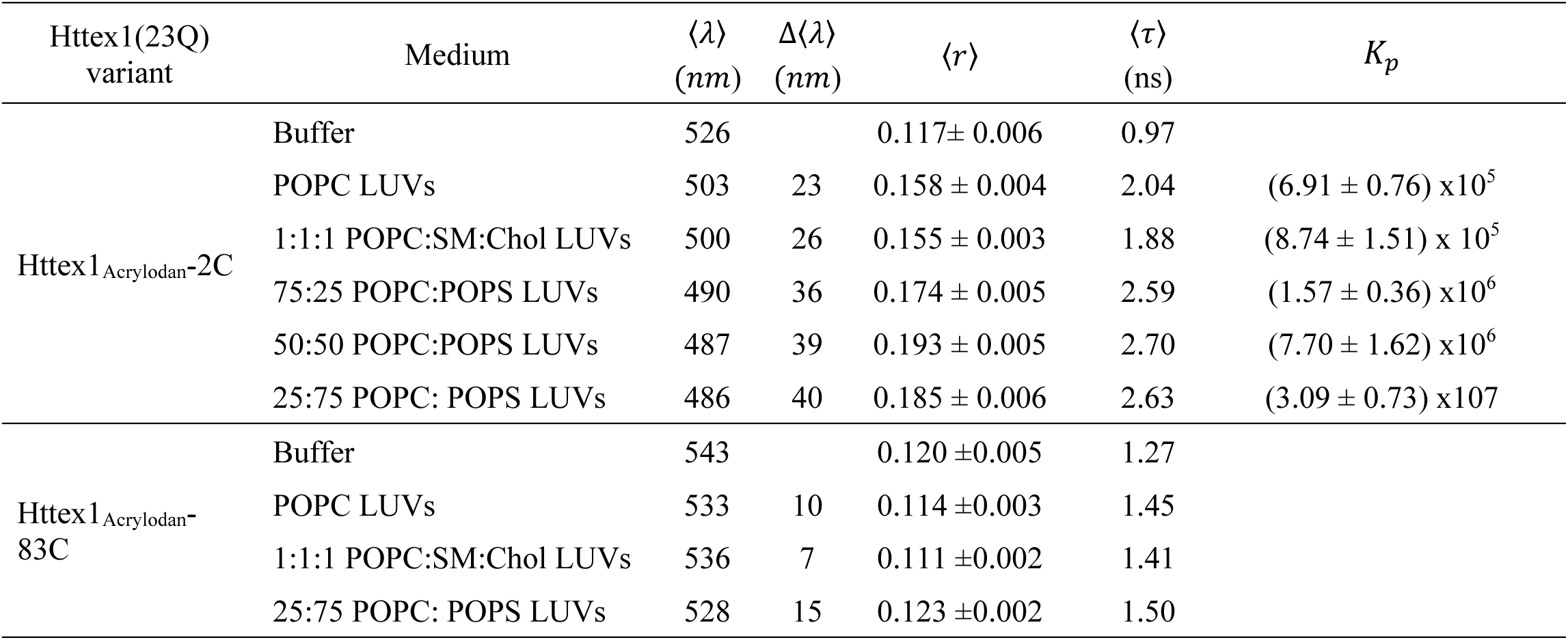
Fluorescence emission properties of Httex1Acrylodan-2C and Httex1Acrylodan-83C variants in solution or in the presence of 1mM LUVs composed of pure POPC, 1:1:1 POPC:SM:Chol and POPC:POPS mixtures. Δ〈𝜆〉 compares the shifts in the fluorescence spectral center-of-mass, 〈𝜆〉, of Httex1(23Q) variants in the absence and in the presence of lipid vesicles. 〈𝑟〉 is the steady-state fluorescence anisotropy and 〈𝜏〉 is the amplitude-weighted mean fluorescence lifetime. The molar partition coefficients, 𝐾*_J_*, were obtained from fitting 〈𝜏〉 data as described in the Materials and Methods section.

| Httex1(23Q)<br>variant | Medium | $\langle\lambda\rangle$<br>(nm) | $\Delta\langle\lambda\rangle$<br>(nm) | $\langle r \rangle$ | $\langle\tau\rangle$<br>(ns) | $K_p$ |
| --- | --- | --- | --- | --- | --- | --- |
| Httex1<sub>Acrylodan</sub>-2C | Buffer | 526 | | 0.117± 0.006 | 0.97 | |
| | POPC LUVs | 503 | 23 | 0.158 ± 0.004 | 2.04 | (6.91 ± 0.76) x10<sup>5</sup> |
| | 1:1:1 POPC:SM:Chol LUVs | 500 | 26 | 0.155 ± 0.003 | 1.88 | (8.74 ± 1.51) x 10<sup>5</sup> |
| | 75:25 POPC:POPS LUVs | 490 | 36 | 0.174 ± 0.005 | 2.59 | (1.57 ± 0.36) x10<sup>6</sup> |
| | 50:50 POPC:POPS LUVs | 487 | 39 | 0.193 ± 0.005 | 2.70 | (7.70 ± 1.62) x10<sup>6</sup> |
| | 25:75 POPC: POPS LUVs | 486 | 40 | 0.185 ± 0.006 | 2.63 | (3.09 ± 0.73) x10<sup>7</sup> |
| Httex1<sub>Acrylodan</sub>-<br>83C | Buffer | 543 | | 0.120 ± 0.005 | 1.27 | |
| | POPC LUVs | 533 | 10 | 0.114 ± 0.003 | 1.45 | |
| | 1:1:1 POPC:SM:Chol LUVs | 536 | 7 | 0.111 ± 0.002 | 1.41 | |
| | 25:75 POPC: POPS LUVs | 528 | 15 | 0.123 ± 0.002 | 1.50 | |

**Table 2.** Typical time-resolved fluorescence anisotropy parameters (rotational correlation times, 𝜙*_A_*, and amplitudes, 𝛽*_A_*) of 0.3 µM Httex1Atto488-2C and Httex1Atto488-83C constructs in aqueous solution (50 mM HEPES, 50mM NaCl, pH 7.4 buffer) and in the presence of 1 mM 25:75 POPC:POPS LUVs (membrane-bound state). The amplitude-weighted mean fluorescence lifetime, 〈𝜏〉, and steady-state fluorescence, 〈𝑟〉, were also determined.

| Httex1(23Q)<br>variant | Medium | $\beta_1$ | $\phi_1$<br>(ns) | $\beta_2$ | $\phi_2$<br>(ns) | $\beta_3$ | $\phi_3$<br>(ns) | $\langle\tau\rangle$<br>(ns) | $\langle r \rangle$ |
| --- | --- | --- | --- | --- | --- | --- | --- | --- | --- |
| Httex1<sub>Atto488-2C</sub> | Buffer | 0.20 | 0.36 | 0.12 | 2.64 | | | 3.35 | $0.076 \pm 0.010$ |
| | 25:75 POPC: POPS LUVs | 0.09 | 0.09 | 0.13 | 0.84 | 0.13 | 30.6 | 3.37 | $0.130 \pm 0.015$ |
| Httex1<sub>Atto488-83C</sub> | Buffer | 0.23 | 0.31 | 0.13 | 1.88 | | | 3.88 | $0.055 \pm 0.002$ |
| | 25:75 POPC: POPS LUVs | 0.24 | 0.22 | 0.15 | 1.40 | | | 3.97 | $0.059 \pm 0.001$ |

### Lipid composition modulates the N-Terminal membrane-interaction of Httex1(23Q), while the C-Terminal remains solvent-exposed

We next sought to evaluate the contribution of distinct membrane lipid compositions and properties to Httex1(23Q)-membrane interaction. Specifically, we focused on probing the hydrophobic/ electrostatic effects and also the impact of membrane phase separation on Httex1(23Q) membrane binding. The fluorescence properties of both Httex1Acrylodan-2C and Httex1Acrylodan-83C constructs were examined upon adding large unilamellar vesicles (LUVs) composed of pure 1-palmitoyl-2-oleoyl-*sn*-glycero-3-phosphocholine (POPC); 1:1:1 POPC:SM:Chol ternary mixture; and finally POPC:POPS mixtures (Figure 2 and Table 1). Considering that acrylodan coupled to both Cys is largely solvent exposed in solution, both a blue-shifted emission and a fluorescence increase will directly report on membranes binding of the fluorescently-labeled protein, as the dye experiences a more hydrophobic environment at the lipid bilayer interface.

**Figure 2:**
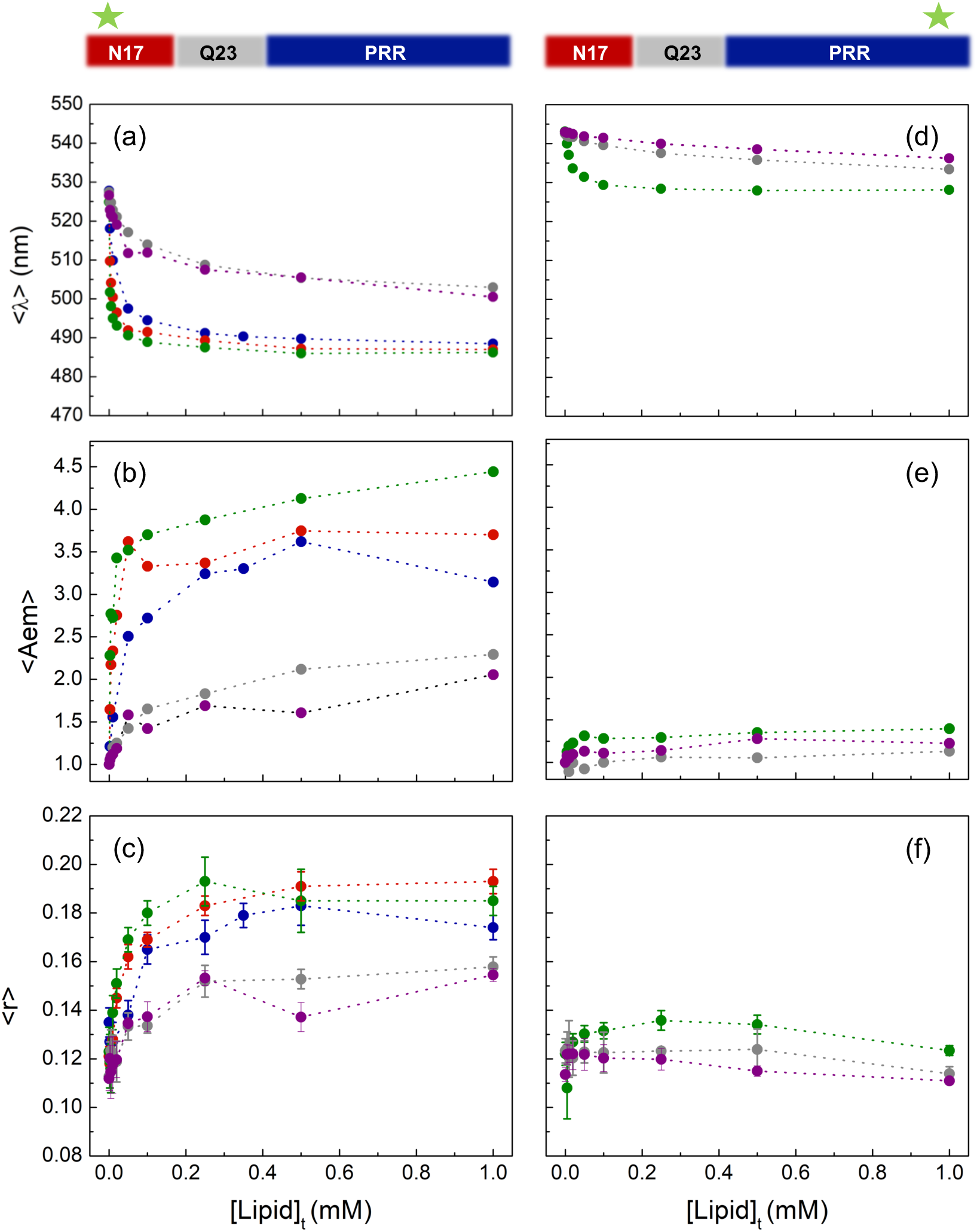
Solvent accessibility of N- and C-termini of Httex1(23Q) upon membrane interaction. Fluorescence emission properties of 0.6 µM (a-c) Httex1Acrylodan-2C and (d-f) Httex1Acrylodan-83C in the presence of LUVs. Changes in the (a and d) fluorescence spectral center-of-mass, (b and e) variation of integrated area, and (c and f) steady-state anisotropy as function of total lipid concentration (ranging from 2.5µM to 1mM). The aqueous solution was 50 mM HEPES, 50mM NaCl, pH 7.4 buffer and LUVs were prepared with pure POPC (grey), 1:1:1 POPC:SM:Chol (purple), 75:25 POPC:POPS (blue), 50:50 POPC:POPS (red) and 25:75 POPC:POPS (green). Lines are just a guide to the eye.

We initially characterized the Httex1Acrylodan-2C variant that maps the N17 segment, since previous studies have indicated that this domain anchors itself to lipid membranes and undergoes a structural transition into an amphipathic α-helix (Atwal et al. 2007; Michalek et al. 2013; Adegbuyiro et al. 2017; Tao et al. 2019). Firstly, its interaction with zwitterionic POPC LUVs was accessed (Figure 1b). Upon increasing the lipid concentration, the fluorescence emission of Httex1Acrylodan-2C underwents a progressive blue-shifted emission (Δ〈𝜆〉*_!"#."_*= 23 nm with 1mM POPC LUVs) and a concomitant increase in the fluorescence intensity (the integrated area under the emission spectrum, 〈𝐴*_/0_*〉, increased 2.3-fold) (Figure 2a, b and Table 1). These results clearly show that acrylodan attached to the N-terminal N17 segment gradually senses a more hydrophobic environment, indicating that this region binds to fluid and zwitterionic lipid membranes. In parallel, the steady- state fluorescence anisotropy of Httex1Acrylodan-2C also enhanced (〈𝑟〉_2C-PC_= 0.158 ±0.004 with 1mM LUVs, Figure 2c and Table 1), reflecting a restricted rotational motion. Overall, these findings reveal that Httex1(23Q)-lipid interaction holds a relevant hydrophobic component.

We also addressed the Httex1Acrylodan-2C association with raft-mimicking lipid membranes. The previous experiments were carried out with 1:1:1 POPC:SM:Chol LUVs with co-existing liquid-disordered (*ld*) and liquid-ordered (*lo*) phases (de Almeida et al. 2003). Globally, the data were comparable to the results obtained for fluid POPC membranes. Upon adding POPC LUVs enriched in Chol and SM, the fluorescence emission again shifted towards shorter wavelengths (Δ〈𝜆〉*_!"#(1&23_*= 26 nm with 1mM 1:1:1 POPC:SM:Chol LUVs), along with an increase in both fluorescence intensity (〈𝐴*_/0_*〉 increased 2.1-fold) and steady-state fluorescence anisotropy (〈𝑟〉_2C-rafts_= 0.155 ±0.003) (Figure 2a-c and Table 1). Altogether, these data indicate that the presence of *lo*-phase did not significantly alter the Httex1(23Q) membrane-interaction, and so Httex1(23Q) seems to interact to a similar extent with both *lo* and *ld* phases.

The electrostatic contribution for Httex1(23Q)-lipid interaction was then assessed using anionic POPS-containing LUVs. The magnitude of changes for all fluorescence parameters obtained for Httex1Acrylodan-2C in the presence of POPC LUVs with 25, 50 and 75 mol% of POPS were higher than for pure POPC and 1:1:1 POPC:SM:Chol vesicles. Upon increasing the concentration of POPC:POPS LUVs, its fluorescence spectra exhibited a more pronounced blue-shift emission (〈𝜆〉*_!"#!4.5_*= 36 nm, 〈𝜆〉*_!"#46.5_*= 39 nm and 〈𝜆〉*_!"#74.5_*= 40 nm with 1mM LUVs), together with a significant increase in fluorescence intensity (Figure 2a, b and Table 1). Moreover, the steady-state fluorescence anisotropy also remarkably enhanced from 〈𝑟〉_2C-buffer_= 0.117±0.009 in buffer to 〈𝑟〉*_!"#!4.5_*= 0.174 ±0.005, 〈𝑟〉*_!"#46.5_*= 0.193 ±0.005 and 〈𝜆〉*_!"#74.5_*= 0.185 ±0.006 with 1mM LUVs (Figure 2c and Table 1). These results reveal that Httex1Acrylodan-2C at anionic lipid surfaces experiences a more hydrophobic environment and a slower rotational mobility (lower 〈𝜆〉 and higher 〈𝑟〉 for POPS-containing vesicles, respectively), supporting that the N-terminal region is more deeply inserted into negatively charged membranes than in pure

Lastly, we examined the interface properties of the C-terminal PRR in the Httex1(23Q) membrane-bound state. The fluorescence properties of Httex1Acrylodan-83C were evaluated in the presence of LUVs composed of pure POPC, 1:1:1 POPC:SM:Chol and 25:75 POPC:POPS (higher anionic content, Figure 2d-f). At variance with the data obtained for Httex1Acrylodan-2C, the emission spectra of Httex1Acrylodan-83C variant showed no significant spectral changes upon adding LUVs (Δ〈𝜆〉*_)*"#."_*= 10 nm, Δ〈𝜆〉*_)*"#81&23_*= 7 nm, and 〈𝜆〉*_)*"#74.5_*= 15 nm with 1 mM LUVs, Figure 2d) and retained its fluorescence intensity (Figure 2e and Table 1), regardless of the lipid composition used. The fluorescence intensity decays for this fluorescently labeled C-terminal construct were well described by similar lifetime components/amplitudes to those in solution state, implicating negligible variations of their amplitude-weighted mean fluorescence lifetime upon varying the lipid concentration (data not shown). The steady-state fluorescence anisotropy also remained mainly unaltered even at higher lipid concentrations (Figure 2f). Altogether, these findings strongly indicate that the C-terminal PRR (83C) remains fully exposed to the aqueous solution and does not sense the hydrophobic membrane surface upon Httex1(23Q) membrane-binding. Moreover, the membrane physical properties- such as the presence of anionic headgroups or lipid phase separation- do not seem to influence the C-terminal solvent accessibility in the Httex1(23Q) membrane-bound state.

### The strength of Httex1(23Q) association with lipid membranes

To obtain a quantitative insight into the role of charge effects and membrane phase separation in Httex1(23Q)-lipid interaction, the mole-fraction partition coefficients, 𝐾_P_, were determined from the amplitude-weighted mean fluorescence lifetime, 〈𝜏〉, data obtained for Httex1Acrylodan-2C (the N-terminal acrylodan-labeled construct). The Httex1(23Q) membrane-binding produced a hyperbolic variation in 〈𝜏〉 with the lipid concentration, consistent with a non-cooperative partitioning model (Figure 3). A detailed analysis of the fluorescence decays of Httex1Acrylodan-2C revealed markedly distinct profiles in solution and in the presence of LUVs (as illustrated Table S1 for 25 mol% POPS). At low lipid concentrations (high free fraction), the decays required four exponential components, similar to buffer. In contrast, as the lipid concentration increased, the fast component disappeared and the decays were well fitted by a sum of three exponentials, with a dominant long-lived component (Table S1). Membrane binding of Httex1Acrylodan-2C increased its 〈𝜏〉 by 2.1-fold for a zwitterionic composition (〈𝜏〉*_!"#$%&&’(_* = 0.97 ns in buffer to 〈𝜏〉*_!"#."_*= 2.04 ns with 1mM POPC LUVs) and 1.9-fold for phase-separated membranes (〈𝜏〉*_!"#81&23_* = 1.88 ns with 1mM POPC:SM:Chol LUVs). An even higher variation in 〈𝜏〉 was obtained for POPS-containing LUVs, with an average 2.7-fold increase across 25–75 mol% POPS (〈𝜏〉*_!"#!4.5_*= 2.59 ns, 〈𝜏〉*_!"#46.5_* = 2.70 ns and 〈𝜏〉*_!"#74.5_* =2.63 ns) (Table 1). These results indicate that the strong self-quenching of acrylodan at 2C in buffer solution is relieved upon membrane binding, leading to a progressive increase in 〈𝜏〉 as Httex1(23Q) partitions into lipid vesicles and becomes incorporated into a less polar environment. These changes are consistent with prior work using other solvatochromic dyes (such as prodan closely related with acrylodan), where more hydrophobic environments result into a fluorescence lifetime increase (Al-Hassan and Khanfer 1998).

**Figure 3:**
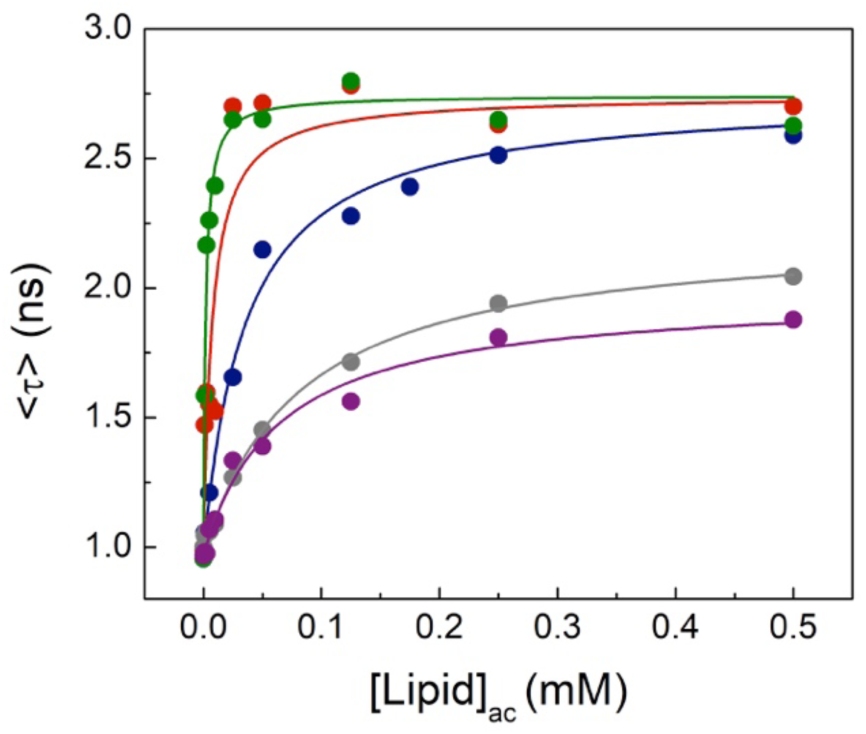
The molar membrane-partition coefficients (𝐾*_J_*) of Httex1(23Q). Variations on the mean fluorescence lifetime of Httex1Acrylodan-2C as function of the accessible lipid concentration, using POPC (grey), 1:1:1 POPC:SM:Chol (purple), 75:25 POPC:POPS (blue), 50:50 POPC:POPS (red) and 25:75 POPC:POPS (green) LUVs. The solid lines represent the best fits of Eq. 5 to the experimental data.

The 𝐾_P_ values for each lipid composition were then retrieved from the variations of 〈𝜏〉 as function of accessible lipid concentration ([Lipid]*_19_*) by fitting EQ. 5 (Figure 3), assuming diluted regime conditions as detailed in Materials and Methods. Importantly, 〈𝜏〉 is unaffected by variations in fluorophore concentration (absolute parameter) and inner filter effects. We performed individual analysis for both pure POPC and 1:1:1 POPC:SM:Chol ternary lipid mixture and the binding energetics were similar for both lipid compositions (𝐾_P-PC_= (6.91 ± 0.76) x10^5^ and 𝐾_P-Rafts_ = (8.74 ± 1.51) x 10^5^ (Table 1)). A global analysis was implemented for POPS:POPS LUVs prepared with variable anionic phospholipid content, since 〈τ〉 converged to a similar plateau at high lipid concentrations. A more marked effect was detected for anionic lipid vesicles, where 𝐾_P_ values increased 2.3 -, 11.1- and 44.7-fold upon adding 25, 50 and 75 mol% of POPS to POPC LUVs (𝐾_P-25PS_= (1.57 ± 0.36) x10^6^, 𝐾_P-50PS_= (7.70 ± 1.62) x10^6^ and 𝐾_P-75PS_= (3.09 ± 0.73) x10^7^ (Table 1)). Overall, these findings highlight that although Httex1(23Q) can associate with both fluid and liquid-ordered membranes, electrostatic effects play the dominant role in Httex1(23Q) lipid binding.

### Distinct conformational dynamics at the N- and C-termini of Httex1(23Q) membrane-bound state

The conformational flexibility of the N-terminal N17 and C-terminal PRR segments of Httex1(23Q) was directly assessed by performing time-resolved fluorescence anisotropy measurements of Httex1(23Q)-A2C or -A83C labeled with Atto 488 maleimide (hereafter termed Httex1Atto488-2C and Httex1Atto488-83C, respectively), both in its free and lipid-bound states. Here, Atto 488 dye was selected due to its general insensitivity to solvent polarity and relatively longer fluorescence lifetime that allows to monitor the rotational dynamics of the fluorescently-labeled protein. Fluorescence correlation spectroscopy (FCS) experiments were also used to check for protein aggregation in aqueous solution and to further validate membrane-binding of both constructs.

We initially characterized the hydrodynamic properties of Httex1(23Q) in aqueous solution by FCS. Here, their translational diffusion times, 𝜏, of both Httex1Atto488-2C and Httex1Atto488-83C were determined in aqueous solution. Remarkably, the diffusion of aggregates/oligomers during the FCS experiments were not detected, which would account for spikes in the fluorescence time traces due to the diffusion of multi-labeled species and with longer diffusion times. The autocorrelation curves were well described by a one-component diffusion model (single diffusing species) and the recovered diffused times were similar for both variants (with comparable diffusion coefficients, 𝐷_2C-buffer_= 6.57 x 10^-11^ m^2^ s^-1^ and 𝐷_83C-buffer_= 6.30 m^2^ s^-1^). We further determined the hydrodynamic radius, 𝑟*_;_*, that reports on size and the compactness of the polypeptide chain (𝑟*_;_*=32 Å for Httex1Atto488-2C and Httex1Atto488-83C, respectively, assuming Stokes-Einstein equation). Particularly, these 𝑟*_;_* values are close to the predicted for a disordered protein (25.5 Å, considering the scaling law 𝑟*^<:=^* = 3.128𝑁*^6.-?!^*(Dudás and Bodor 2019) for a polymer length, 𝑁), and further indicate that Httex1(23Q) does not adopt an overall collapsed ensemble in solution.

To exclude artifacts from Cys mutations and the fluorescent-labeling in Httex1(23Q)-membrane binding, FCS measurements were also performed with Httex1Atto488-2C and Httex1Atto488-83C (using 10 nM protein concentration) in the presence of 25:75 POPC:POPS LUVs. We note that FCS data should be independent of the labeling site. The fluorescence time traces showed no spikes, indicating that POPS-containing LUVs did not induce Httex1(23Q) aggregation. Moreover, the autocorrelation curves of both Httex1Atto488-2C and Httex1Atto488-83C shifted towards longer time scales upon adding 1mM 25:75 POPC:POPS LUVs (Figure S1- for Httex1Atto488-83C), showing that both constructs effectively bind to anionic lipid vesicles.

We then characterized the general fluorescence properties of both Httex1Atto488-2C and Httex1Atto488-83C constructs in buffer and with 1 mM 25:75 POPC:POPS LUVs. Under this last condition (high anionic lipid content and lipid concentration), the previous acrylodan data indicate that the majority of the Httex1(23Q) population is bound to LUVs (membrane-bound fraction ∼ 0.98 retrieved from 𝐾_P_) and that its fluorescently-labeled residue 2C (within N17 segment) becomes more deeply inserted into the lipid bilayer compared to zwitterionic/phase separated membranes. As expected, for the Atto 488 probe, no major differences were recorded in the mean fluorescence lifetime and the fluorescence emission spectra of Atto 488 at either Cys labeling site upon membrane association. At variance, the steady-state fluorescence anisotropy, 〈𝑟〉, of Httex1Atto488-2C increased upon adding anionic LUVs (from 〈𝑟〉 = 0.076 ± 0.010 in solution to 〈𝑟〉 = 0.130 ± 0.015 with 1mM 25:75 POPC:POPS LUVs), confirming the membrane-anchoring of its N17 segment. To obtain further information on the different time scales of the rotational motions contributing to the protein dynamics, fluorescence anisotropy decays were obtained for both Httex1Atto488-2C and Httex1Atto488-83C constructs.

In aqueous solution (free state), the time-resolved fluorescence anisotropy of Httex1Atto488-2C and Httex1Atto488-83C rapidly decayed to zero, confirming the absence of large protein aggregates in buffer. Moreover, the anisotropy decays exhibited a typical two-exponential depolarization kinetics that likely arises from the sub-nanosecond local rotational mobility of Atto 488 (fast rotational correlation time, 𝜙,) and the segmental motion (slow rotational correlation time, 𝜙*_!_*). Remarkably, a similar fast correlation time was recovered for both labeling positions (𝜙*_,,!"#$%&&’(_* ≈ 0.36 ns and 𝜙*_,,)*"#$%&&’(_* ≈ 0.31 ns), accounting for > 60% of the depolarization. In contrast, the segmental fluctuations were slightly distinct between the two termini (𝜙*_!,!"#$%&&’(_* ≈ 2.6 ns and 𝜙*_!,)*"#$%&&’(_* ≈ 1.9 ns). This time-range of correlation times has been assigned in IDPs to the backbone motion of the polypeptide chain (Majumdar and Mukhopadhyay 2018; Bhasne et al. 2020; Das et al. 2021). Therefore, the data support that the C-terminal (83C) displays a faster segmental flexibility as typical of IDPs, whereas the N-terminal (A2C) shows slower dynamics, consistent with a slightly more locally compacted chain in solution.

Upon addition of 1mM POPS-containing LUVs (membrane-bound state), the fluorescence anisotropy decays for Httex1Atto488-2C and Httex1Atto488-83C constructs diverged significantly. For Httex1Atto488-2C (N-terminal), the anisotropy decayed much slower than in solution and was adequately described by a tri-exponential decay function. Specifically, two short rotational correlation times (𝜙*_,,!"#74.5_*≈ 0.09 ns and 𝜙*_!,!"#74.5_* ≈ 0.84 ns) and a much longer component (𝜙*_*,!"#74.5_* ∼30 ns) were obtained. These data show the absence of complete immobilization (while a residual anisotropy was not detected at longer times). Moreover, the third long component indicate the slowdown of the rotational motion around the N-terminal in the presence of anionic liposomes. In opposite, the anisotropy decays for Httex1Atto488-83C remained nearly identical to the solution state and retained similar timescales for both rotational correlation times (𝜙*_,,)*"#74.5_* ≈ 0.22 ns and 𝜙*_!,)*"#74.5_* ≈ 1.4 ns). This lack of change highlights that the C-terminal does not directly interact with lipid vesicles and holds highly dynamic features in the Httex1(23Q) membrane-bound state.

## Discussion

The dysregulated association of Htt with multiple membranous organelles (such as ER, Golgi apparatus and mitochondria) and synaptic vesicles (DiFiglia et al. 1995; Velier et al. 1998; Li et al. 2000; Panov et al. 2002; Choo et al. 2004; Atwal et al. 2007; Rockabrand et al. 2007; Ueda et al. 2016), each with distinct biophysical membrane properties, has been implicated in HD pathogenesis. The complex interplay between Httex1 and cellular membranes represents a critical aspect of HD, and its earliest interaction stages could provide a promising therapeutic target for halting HD progression. Previous studies on Httex1-lipid interaction have primarily examined highly aggregation-prone conditions (often employing circular dichroism or Thioflavin T assays) and a restricted range of lipid compositions or complex mixtures (including TLBE). The present work provides a quantitative characterization of the initial steps of monomeric Httex1(23Q) association with lipid membranes, with particular emphasis on clarifying the contributions of each flanking polyQ regions - N-terminal N17 segment and C-terminal PRR. Using site-specific fluorescence labeling with two different fluorescent reporters (acrylodan and Atto488 dyes) and a systematic variation of the lipid composition, we probed the conformational dynamics and membrane interaction of both termini of Httex1(23Q) across diverse lipid environments, including zwitterionic, anionic, and phase-separated membranes.

The fluorescence emission spectra obtained for the acrylodan-labeled Httex1(Q23) variants (Httex1Acrylodan-2C and Httex1Acrylodan-83C) confirmed that both N- and C-terminal regions of monomeric Httex1(23Q) experience polar environments in the free state. A similar range of emission maxima was reported for acrylodan-labeled IDPs in aqueous solution, namely for 𝛼- synuclein and Importinβ-binding domain of Importinα (IBB) (Arya et al. 2018; Chowdhury et al. 2019). However, a closer inspection of the spectral center of mass and mean fluorescence lifetime of acrylodan covalently-linked at the 2C and 83C positions indicate variations between both termini (Table 1), underlining distinct local conformational properties within N17 and PRR domains. Specifically, acrylodan coupled to residue 2C exhibited a shorter 〈𝜆〉 and complex fluorescence decay, indicating that the N17 segment samples a less polar and more heterogeneous environment. These findings are also aligned with the slower segmental motion retrieved for Httex1Atto488-2C. Globally, our data for monomeric Httex1(23Q) in buffer are consistent with the “tadpole-like” topology proposed from NMR and smFRET experiments (Warner et al. 2017; Newcombe et al. 2018), where N17 adopts a more compact topology and PRR is highly solvent-exposed and flexible.

Upon Httex1(23Q) membrane association, the N17 segment exhibited hallmarks of a membrane-binding motif. Here, acrylodan labeling at residue 2C, which is positioned adjacent to the amphipathic 𝛼-helix (extended from residues 3–13) (Tao et al. 2019), showed a gradual blue-shifted emission and enhanced fluorescence intensity with increasing lipid concentrations, consistent with its membrane anchoring (within a more hydrophobic environment) (Figure 2a, b). Steady-state anisotropy measurements further show the membrane association (Figure 2c) and the restricted rotational mobility of Httex1Atto488-2C (Figure 4a). Collectively, these results are in agreement with the disorder-to-helix transition of N17 upon membrane interaction reported by Michalek et al (Michalek et al. 2013) and Tao et al. (Tao et al. 2019) using other biophysical methods and here mainly centered in anionic lipid membranes.

**Figure 4.**
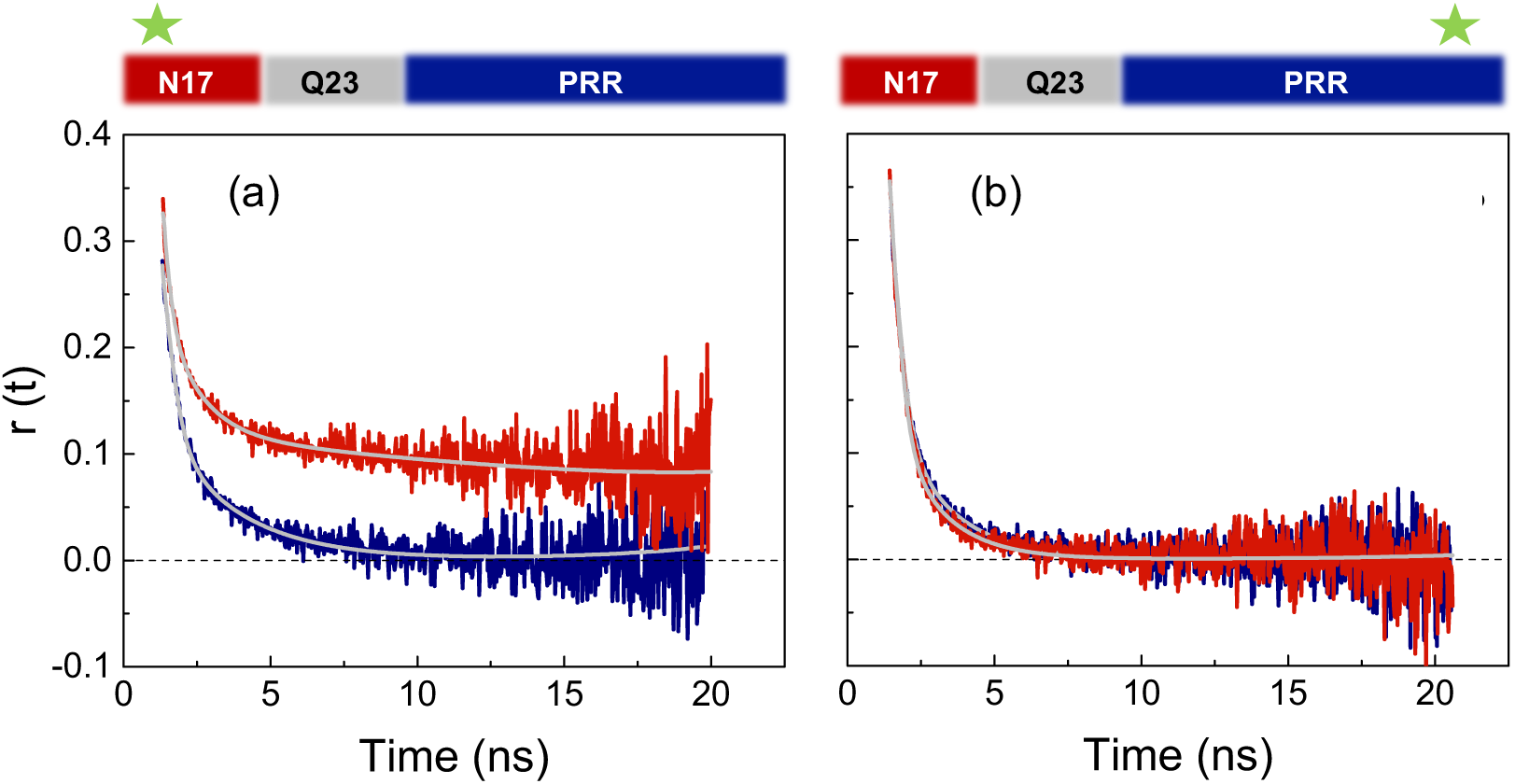
Comparison of the typical fluorescence anisotropy decays of 0.3 µM (a) Httex1Atto488- 2C and (b) Httex1Atto488-83C in the free (50 mM HEPES, 50mM NaCl, pH 7.4 buffer) and the membrane-bound state (1mM 25:75 POPC:POPS LUVs). All time-resolved anisotropy data were adequately fitted with a bi-exponential decay function, with exception of Httex1Atto488-2C in the membrane-bound form that required tri-exponentials. The fits are shown by the solid grey lines.

Importantly, our systematic lipid screen reveals that electrostatics play a dominant but not exclusive role, in controlling Httex1-membrane interaction. The anionic POPS lipid markedly enhanced Httex1(23Q) affinity, with membrane-partition coefficients increasing up to 44-fold (for 75 mol% POPS) compared with zwitterionic POPC vesicles (Table 1). Our data further indicate that N17 segment inserts deeper into the lipid bilayer (compared with zwitterionic fluid and phase separated lipid membranes), and exhibits reduced rotational mobility. These results are in line with previous reports showing that negatively charged lipids accelerate Httex1 aggregation/fibrillation, and that lipid headgroup charge directly modulates both the kinetics and morphology of Httex1 aggregates (Beasley et al. 2021). In this context, the positive net charge of N17 (comprising 3 lysines and 2 glutamic acids) seems crucial for promoting favorable electrostatic contacts with anionic lipids, thereby stabilizing the Httex1(23Q) –membrane interface.

Our acrylodan fluorescence data also indicate that Httex1(23Q) associates to a similar extent with both fluid POPC vesicles and raft-mimicking membranes composed of 1:1:1 POPC:SM:Chol (Table 1). Interestingly, the inclusion of SM and Chol does not appear to reduce the Httex1(23Q) membrane association, at variance with previous reports using TLBE enriched in SM (Chaibva et al. 2018) or Chol (Gao et al. 2016), or even POPC with Chol (Stonebraker et al. 2023). This discrepancy may reflect the complexity of the lipid extracts (such as TLBE) and the sensitivity of the techniques employed.

In complete contrast to N17 segment, minimal changes in the acrylodan fluorescence properties or rotational dynamics of Atto488 were detected for the Httex1(23Q)-83C construct (N-terminal PRR) upon its membrane association. These observations are consistent with previous reports showing that the PRR (with a negative net charge) remains largely solvent-exposed in the presence of anionic lipid vesicles (Tao et al. 2019). Our data further demonstrate that this lack of membrane direct association is independent of the lipid composition, and remarkably the PRR segment retains its high conformational flexibility even in the membrane-bound state. Similar behavior is also displayed by the C-terminal of α-synuclein upon membrane binding (Fusco et al. 2014; Bhasne et al. 2020).

In summary, the present work reveals that the initial membrane recruitment of Httex1 (23Q) is driven by both electrostatic and hydrophobic interactions, with distinct contributions from the N17 segment and PRR. Here, we show that anionic POPS-containing membranes drive deeper insertion of the N-terminal N17 into the lipid bilayer, inducing restricted conformational dynamics. In contrast, the PRR retains its solution disordered and dynamic features, regardless of the lipid composition (Figure 5). These distinct membrane-binding behaviors provide insight into the broad subcellular distribution of Httex1 fragments across organelles with diverse lipid environments and also provide mechanistic details into how membrane composition could modulate aggregation propensity in HD. Future studies should investigate how post-translational modifications within N17, such as phosphorylation or acetylation, influence its conformational dynamics. Additionally, exploring the consequences of PRR solvent-exposure, particularly in its ability to interact with other proteins upon membrane binding, will be crucial to untangling its role in Httex1 aggregation. Integrating these biophysical insights will be essential for developing therapeutic strategies aimed to control the early pathogenic interactions of Httex1 with lipid membranes, potentially leading to novel therapeutic approaches for HD.

**Figure 5.**
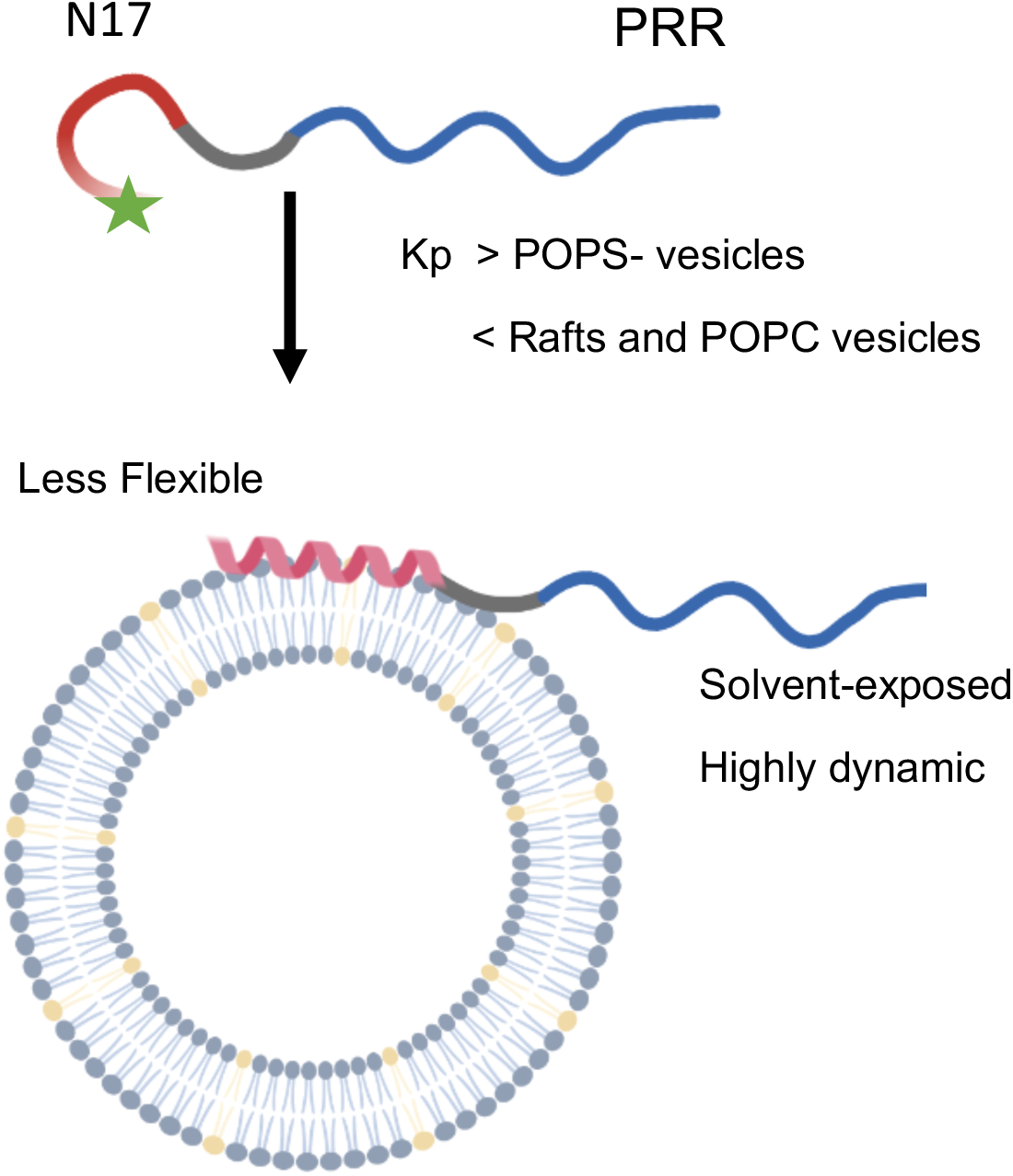
Proposed model for the membrane binding of monomeric Httex1(23Q). Our data reveal a favored partitioning of Httex1(23Q) towards POPS-containing LUVs. Nevertheless, Httex1-lipid interaction is not solely controlled by electrostatic effects (still binds to POPC and 1:1:1 POPC:SM:Chol LUVs). In particular, the N-terminal N17 region undergoes a deep membrane insertion and adopts a less flexible conformation, whereas the C-terminal PRR retains its dynamic and solvent-exposed features, regardless of the lipid composition.

## MATERIALS AND METHODS

### Materials

Champion™ pET SUMO expression system and BL21(DE3) One Shot® chemically competent *E. coli* cells were purchased from Invitrogen. QuikChange II site-directed mutagenesis kit was from Agilent. The primers were synthesized by STAB VIDA. Luria Broth (Miller’s LB Broth), isopropyl β-D-1-thiogalactopyranoside (IPTG), dithiothreitol (DTT) and kanamycin (Km) were purchased from NZYTech. The cOmplete EDTA-free protease inhibitor cocktail tablets were from Roche. Phenylmethylsulfonyl fluoride (PMSF) was obtained from PanReac AppliChem. Amicon Ultra-15 centrifugal filters were purchased from Millipore. 0.44 μm syringe filters (low protein binding) were from Labbox. The Pierce™ BCA protein assay kit and Slide-A-Lyzer dialysis cassettes were from Thermo Fisher. The 5-mL HisTrap FF, 5-mL HiTrap desalting, and Superdex 75 10/300 GL columns were obtained from Cytiva.

The POPC, POPS and brain SM (Brain, Porcine) were from Avanti Polar Lipids, while Chol was from Sigma. The 6-acryloyl-2-dimethylaminonaphthalene (Acrylodan) was obtained from Invitrogen and Atto 488 maleimide (Atto 488) was from Sigma.

Chemicals for buffer preparation used in protein purification, labeling, and the fluorescence experiments were obtained from Thermo Fisher. Buffers were prepared with Milli-Q water (18.2 MΩ·cm) and further filtered with 0.2 μm membrane filters (Sartorius).

### Plasmid constructs

Httex1(23Q) cDNA was amplified by PCR from the plasmid pTWIN1-His6-Ssp-Httex1-23Q (gift from Hilal Lashuel - Addgene plasmid #84349) (Vieweg et al. 2016) using the forward primer 5’- GCCACCCTGGAAAAACTGATGA-3’ and the reverse primer 5’- TACGGACGATGCAGCGGTT-3’. The PCR product was then inserted into a pET SUMO vector using TA cloning, as described in the manufacturer’s protocol. This vector allows expression with a N-terminal tag containing a polyhistidine (His6) and a small ubiquitin-related modifier (SUMO) protein. For site-specific fluorophore labeling of Httex1(23Q), a single cysteine at position A2C or A83C was introduced using QuikChange II Site-Directed Mutagenesis. All cloning and mutagenesis were verified by DNA sequencing (STAB VIDA, Portugal). Finally, the pFGET19_Ulp1 plasmid used for SUMO Protease production (the catalytic domain of the ubiquitin-like protease 1, Ulp1403–621) was a gift from Hideo Iwai (Addgene plasmid # 64697) (Guerrero et al. 2015).

### Httex1 expression and purification under native conditions

BL21(DE3) *E. coli* cells were transformed with pET-SUMO-Httex1(23Q) plasmid (with A2C or A83C mutations) using Km as the selection antibiotic. A single colony was inoculated into 25 mL of LB media with 50 µg/mL Km at 37 °C overnight (O/N) in a shaking incubator. Afterwards, 10 mL of the O/N culture was renewed into 100 mL of LB-Km media and further incubated for 3/4 hours under the same conditions. For protein production, 1L of LB-Km media was set to an OD600 of 0.1 (from the previous culture) and further cultured to OD600 ∼ 0.4 - 0.6 at 37 ⁰C. Protein expression was then induced with 0.6 mM IPTG and cells were grown O/N at 16 °C under continuous shaking. Finally, cells were pelleted by centrifugation (8000 rpm, 4 ⁰C, 10 min) and re-suspended into 60 mL of Buffer Nickel A (50 mM Tris-HCl, 500 mM NaCl, 15 mM imidazole, pH 8) containing cOmplete EDTA-free protease inhibitors and 1 mM PMSF.

Cells were lysed on ice by sonication (using Branson Sonifier 250) and cell debris were removed by centrifugation (17600 x *g*, 4 ⁰C, 1 hour). The supernatant was filtered through 0.44 μm syringe filter (low protein binding) and loaded into a 5-mL HisTrap FF column at 1.5 mL/min using an ÄKTA start chromatography system (Cytiva). The non-specific bound proteins were washed out with Buffer Nickel A at 5 mL/min. The His6-SUMO-Httex1(23Q) fusion protein was then eluted with 100% Buffer Nickel B (50 mM Tris-HCl, 500 mM NaCl, 500 mM imidazole, pH 8) at 1.5 mL/min. Fractions containing the fusion protein were pooled and buffer exchanged to Buffer Nickel C (50 mM Tris-HCl, 150 mM NaCl, 15 mM imidazole, pH 8) in Amicon Ultra-15 with 10 kDa MWCO. To cleave the His6-SUMO tag, the protein was incubated with Ulp1403–621 (SUMO protease, 1:50 v:v) and 1mM DTT during 3 hours at 4 °C in a rotator mixer. The tag-free Httex1(23Q) protein was then purified from the His6-SUMO-tag, fusion protein, and His6-ULP1 by a second 5-mL HisTrap FF column. Here, the Httex1(23Q) protein was collected in the flow- through at 1.5 mL/min. The fractions containing the tag-free protein were pooled and concentrated (∼500 μL) into SEC Buffer (50 mM Tris-HCl, 200 mM NaCl, pH 8.0) using an Amicon Ultra-15 3 kDa MWCO. Final purification was achieved by size exclusion chromatography (SEC) on a Superdex TM 75 10/300 GL column at 0.3 mL/min using an AKTA Purifier 10 (GE Healthcare). The elution of Httex1(23Q) (fusion and tag-free) was analyzed by SDS-PAGE and Western Blot (using a primary mouse anti-Htt MAB5492). Protein concentration was assessed using the Pierce BCA protein assay kit.

### Httex1 site-specific labeling with acrylodan and Atto488 maleimide

The labeling of Httex1(23Q) at positions A2C or A83C with acrylodan or Atto488 maleimide was carried out as previously described (Melo et al. 2016; Melo et al. 2017) with minor modifications. Briefly, freshly purified protein (typically 500–1000 μL, with >100 μM protein) was incubated with 1 mM DTT for 30 min at room temperature (RT). The protein was then loaded into two coupled HiTrap Desalting Columns for exchanging into the labeling buffer (20 mM Tris, 50 mM NaCl, 6 M guanidine hydrochloride (GdmCl), pH 7.4) and remove DTT. The Httex1(23Q) was then incubated with a tenfold molar excess of the dye during 4 h at RT for acrylodan and O/N at 4 °C for Atto488 maleimide, respectively. Finally, the unreacted dye and GdmCl were removed by passing over a set of coupled desalting columns equilibrated in SEC buffer. Fractions containing Atto488- or acrylodan-labeled Httex1(23Q) (at A2C and A83C positions) were aliquoted into 20- 30 μL and flash frozen in liquid nitrogen for final storage at -80 ⁰C. The dyes concentrations were determined spectrophotometrically using molar extinction coefficients of 12 900 M^−1^cm^−1^ at 360 nm (Prendergast et al. 1983) for acrylodan, and 90 000 M^−1^ cm^−1^ at 500 nm for Atto488.

### Ulp1 protease production

The expression and purification of the SUMO protease (Ulp1) was adapted from Guerrero et. al, 2015 method (Guerrero et al. 2015). Briefly, N-terminal His6-tagged Ulp1403–621 protease was expressed from pFGET19_Ulp1 plasmid (Km as selection antibiotic) in *E. coli* BL21(DE3). An isolated colony was picked to create 100 mL LB-Km overnight culture at 37⁰C. 1L of LB-Km culture was then started with an OD600 of 0.1 (from the O/N culture), and the cells were grown at 37⁰C until reaching an OD600nm ∼ 0.5. 1mM IPTG was added for Ulp1 expression at 37 °C during 4 h. Afterwards, cells were harvested as described for Httex1(23Q) and resuspended in Buffer A- Ulp1 (50 mM sodium phosphate, 300 mM NaCl, pH 8.0).

For the Ulp1 purification, cells were lysed and cell debris removed as detailed for Httex1(23Q). A 5-mL HisTrap FF column was initially loaded with the cell extract and then washout out with Buffer A-Ulp1 at 5 mL/min. The Ulp1 was eluted with Buffer B-Ulp1 (50 mM sodium phosphate, 300 mM NaCl, 250 mM imidazole, pH 8.0) using a gradient from 0-100% at 5 mL/min. Fractions containing the pure protease were dialyzed O/N at 4 °C against phosphate buffered saline (PBS) buffer with 50% glycerol and 25 mM DTT. The ULP1 was then aliquoted, flash-frozen and stored at -80 °C.

### Liposome Preparation

LUVs of pure POPC; POPC mixtures containing 25, 50 or 75 mol % of POPS; and finally 1:1:1 POPC:SM:Chol ternary mixture were prepared by the extrusion method (Mayer et al. 1986). The concentrations of the lipid stock solutions were prepared in chloroform UVASOL (Merck). Briefly, multilamellar vesicles (MLVs) in 50 mM HEPES, 50mM NaCl, pH 7.4 buffer were extruded through an Avanti Mini-Extruder system with 50 nm pore polycarbonate membranes (Whatman). The homogeneity of the lipid vesicles was confirmed by Dynamic Light scattering using a Nanosizer ZS (Malvern Instruments).

### Steady-State Fluorescence Spectroscopy

Steady-state fluorescence experiments were conducted in a Fluorolog-3-21 spectrofluorometer (Horiba Jobin Yvon) with double excitation / emission monochromators, and automated polarizers. Measurements were performed in 0.5×0.5 cm quartz cuvettes (Hellma Analytics) at 23 °C. Fluorescence emission spectra and anisotropy of Httex1(23Q) labeled with acrylodan (0.6 µM) or Atto488 (0.6 µM) in solution and upon adding LUVs were recorded with excitation at 370 and 488 nm, respectively. The fluorescence spectral center-of-mass, 〈𝜆〉, (intensity-weighted average emission wavelength) was calculated from the emission spectra according to:

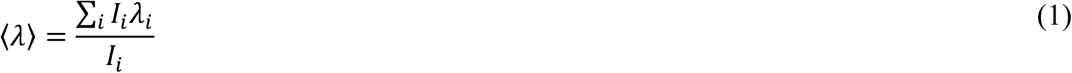

where 𝐼*_A_* is the fluorescence intensity measured at the respective wavelength 𝜆*_A_* (Lopes et al. 2004). The steady-state fluorescence anisotropy, 〈𝑟〉, was determined as (Lakowicz 2006Principles of Fluorescence Spectroscopy):

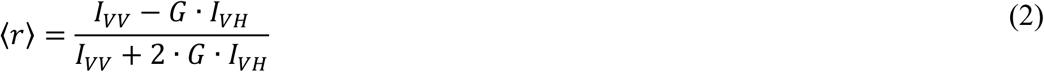

where the intensities 𝐼*_BB_* and 𝐼*_B;_*correspond to the vertically and horizontally polarized emission, using vertically polarized light excitation, respectively. The 𝐺 factor (𝐺 = 𝐼*_;B_*/𝐼*_;;_*, with horizontal excitation) corrects for the transmission efficiency of the monochromator to the polarization of the light. Blank subtraction was always performed to correct for the background signal from buffer and each lipid concentration of LUVs.

### Time-resolved Fluorescence Spectroscopy and decay analysis

Time-resolved fluorescence intensity and anisotropy measurements were performed by the time-correlated single-photon timing technique as previously described (Melo et al. 2013; Scanavachi et al. 2021), Specifically, Httex1(23Q) labeled with acrylodan or Atto488 were excited at 340 nm (with a frequency doubled secondary cavity-dumped dye laser of DCM -Coherent 701-2) and 488 nm (BDS-SM-488FBE pulsed picosecond diode laser from Becker & Hickl), respectively. The emission was recorded as detailed for steady-state fluorescence anisotropy. The fluorescence intensity decays, 𝐼(𝑡), were obtained with the emission polarizer at 54.7° (the magic angle) relative to the vertically polarized excitation beam. For Atto488-labeled Httex1(23Q), anisotropy decays were also recorded; specifically, the parallel and perpendicular polarized decays (𝐼*_BB_*(𝑡) and 𝐼*_B;_*(𝑡), respectively) to the plane of the excitation light were alternatively acquired. The instrument response function (IRF) was obtained from the excitation light scattered by a Ludox solution (silica, colloidal water solution, Aldrich). The IRF and fluorescence intensity decays were acquired in 1024 channels, typically with up to 50 000 and 20 000 counts in the peak channel, respectively. Fluorescence intensity decays, 𝐼(𝑡), were analyzed by a sum of discrete exponential terms:

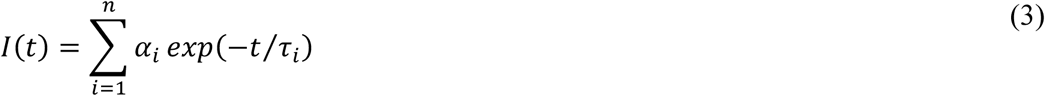

where 𝛼*_A_*is the normalized amplitude and 𝜏*_A_*is the lifetime of the ith decay component of fluorescence. The amplitude-weighted mean fluorescence lifetime, 〈𝜏〉, (proportional to the area under the decay curve or quantum yield), was calculated as:

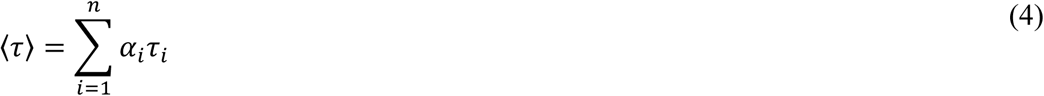

The mole-fraction membrane partition coefficient, 𝐾*_=_*, was obtained by fitting the following equation to the experimental data of 〈𝜏〉 versus [Lipid]ac (under “diluted regime”, the number of moles of water/lipid is much larger than the number of moles of Httex1(23) in each fraction):

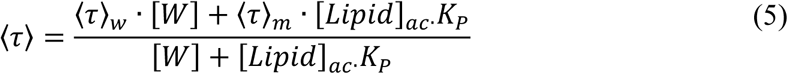

where [𝐿𝑖𝑝𝑖𝑑]*_GH_* and [𝑊] represent the accessible lipid (half of total lipid concentration) and water concentrations, respectively. 〈𝜏〉*_E_* and 〈𝜏〉*_F_* are the mean fluorescence lifetime of the free and in the completely membrane-bound state, respectively.

The fluorescence anisotropy decays, 𝑟(𝑡), were analyzed by global fitting by a sum of discrete exponential terms (Melo et al. 2014a):

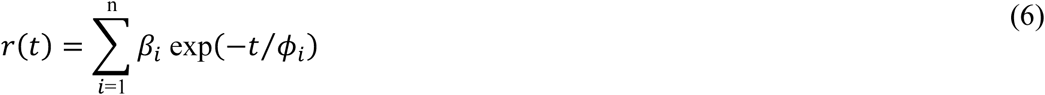

where 𝛽*_A_* and 𝜙*_A_* are the normalized amplitude and the rotational correlation time of the 𝑖th anisotropy decay component, respectively.

All decay analysis were carried out using the TRFA Data Processing Package (version 1.4) from the Scientific Software Technologies Center (Belarusian State University). The goodness of the fits was assessed by the reduced χ2 value (< 1.3) and a random distribution of weighted residuals /autocorrelation plots.

### FCS experiments

FCS measurements were carried out in a home-built confocal setup based on Olympus IX73 Microscope. A Sapphire 488-50 CW laser (Coherent) was used for excitation (∼5 μW). The fluorescence emission was collected by a UPLSAPO 60x/1.2NA water-immersion objective (Olympus) and separated from laser excitation with a ZT488rdc Dichroic Mirror and an ET500lp Long-Pass Filter (Chroma). Photon detection was by fiber (50-μm-diameter aperture; OzOptics) coupled into an APD detector (SPCM-AQRH-14-FC from Excelitas). A digital correlator (Flex03LQ-12; Correlator.com) was used to generate the autocorrelation curves. The autocorrelation curves were fitted to a function for single (free protein) or two-component diffusion (protein upon incubation with LUVs) as previously described (Melo et al. 2011; Melo et al. 2014b).

### Author Contributions

Tânia Sousa: Investigation, Methodology, Formal Analysis; Gonçalo Damas: Investigation, Formal Analysis; Ana Coutinho: Methodology, Formal Analysis; Nuno Bernardes: Methodology; Resources; Ana Azevedo: Methodology, Resources; Manuel Prieto: Methodology, Formal Analysis; Ana M. Melo: Conceptualization, Funding Acquisition, Supervision, Methodology, Formal Analysis, Writing – Review & Editing.

## ACKNOWLEDGMENT

This work was supported by the FCT-Portugal (PTDC/BIA-BFS/30959/2017 and 2022.01454.PTDC grants to Ana M Melo; UID/04565 and LA/P/0140/2020 funding to iBB and i4HB; and UID/00100/2023 to CQE) and Maria de Sousa Prize - 2nd edition funded by Bial Foundation and PT Medical Association to Ana M Melo (Grant 45/2022).

