## Supplementary material for "Lipid Composition Controls the Huntingtin Exon1 Membrane-Association and Differentially Modulates its Flanking Regions Dynamics": Table S1 and Figure S1

**Table S1:** Partitioning of Httex1$_{Acrylodan}$-2C into POPC LUVs with 25 mol% POPS. Typical parameters associated with the time-resolved fluorescence intensity of Httex1$_{Acrylodan}$-2C with increasing lipid concentrations.

| [L] (mM) | $\alpha_1$ | $\tau_1$ (ns) | $\alpha_2$ | $\tau_2$ (ns) | $\alpha_3$ | $\tau_3$ (ns) | $\alpha_4$ | $\tau_4$ (ns) | $\chi^2$ | $\bar{\tau}$ (ns) |
|---|---|---|---|---|---|---|---|---|---|---|
| **0** | 0.33 | 0.10 | 0.38 | 0.51 | 0.23 | 1.7 | 0.06 | 4.3 | 1.08 | 0.87 |
| **0.005** | 0.34 | 0.12 | 0.33 | 0.56 | 0.27 | 2.0 | 0.06 | 4.9 | 1.06 | 1.0 |
| **0.01** | 0.50 | 0.05 | 0.22 | 0.50 | 0.22 | 2.1 | 0.06 | 5.0 | 1.25 | 0.89 |
| **0.02** | - | - | 0.39 | 0.26 | 0.38 | 1,7 | 0.22 | 4.1 | 1.21 | 1.7 |
| **0.05** | - | - | 0.34 | 0.23 | 0.36 | 1.7 | 0.30 | 4.0 | 1.16 | 1.9 |
| **0.10** | - | - | 0.25 | 0.26 | 0.37 | 1.9 | 0.38 | 4.0 | 1.17 | 2.3 |
| **0.25** | - | - | 0.23 | 0.18 | 0.35 | 2.0 | 0.42 | 4.0 | 1.05 | 2.4 |
| **0.50** | - | - | 0.19 | 0.14 | 0.35 | 2.0 | 0.47 | 3.9 | 1.12 | 2.6 |
| **1.0** | - | - | 0.15 | 0.13 | 0.42 | 2.3 | 0.43 | 4.0 | 0.99 | 2.7 |

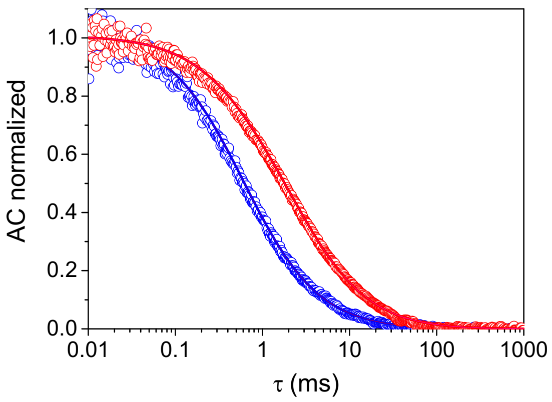


Figure S1: A- Normalized auto-correlation curves of Httex1$_{Atto488}$-83C in the absence (blue) and in the presence of 1 mM POPC:POPS 25:75 LUVs (red). FCS shows no Httex1-23Q aggregation in solution and Httex1-23Q (with a Cys at 83 residue) binds to anionic lipid membranes.
